# The effects and mechanism of peiminine-induced apoptosis in human hepatocellular carcinoma HepG2 cells

**DOI:** 10.1101/377069

**Authors:** Xu Chao, Guoquan Wang, Yuping Tang, Changhu Dong, Hong Li, Bin Wang, Jieqiong Wu, Jiarong Zhao

## Abstract

Peiminine is a compound that is isolated from *Bolbostemma paniculatum* (Maxim) Franquet (Cucurbitaceae family), which has demonstrated antitumor activities. Its precise molecular mechanisms underlying antitumor activity remain elusive. In this study, peiminine-induced apoptosis towards human hepatocellular carcinoma and its molecular mechanisms were investigated. MTT assay was employed to assess anticancer effects of peiminine at concentrations of 2, 4, 6, 8, 10, 12, and 14 μg/ml after 24, 48, or 72 h. Nuclear staining and flow cytometry were carried out to further assess apoptosis. Mitochondrial membrane potential evaluation and Western blot analysis were performed to investigate the mechanism of peiminine-induced apoptosis. Peiminine reduced the viability of HepG2 cells in a time- and dose-dependent manner and had an IC_50_ of 4.58 μg/mL at 24h. Flow cytometry assessment indicated that peiminine markedly increased the cell number of apoptotic cells and the mitochondrial membrane potential dose-dependently in HepG2 cells. The results of Western blotting showed the expression of Bcl-2, procaspase-3, procaspase-8, procaspase-9, and PARP_1_ decreased in HepG2 cells treated with peiminine, while the expression of Bax, caspase-3, caspase-8, caspase-9, and cleaved PARP_1_ increased. The result suggest taht peiminine can induce apoptosis in human hepatocellular carcinoma HepG2 cells through both extrinsic and intrinsic apoptotic pathways.

## Introduction

Hepatocellular carcinoma (HCC) ranks third among malignancies related cancer-related deaths, and annually occurs in approximately 600,000 individuals worldwide[1]. Although significant advances in frontline cancer research and chemotherapy have been made in treating HCC, many of the proposed drugs cause potent toxic adverse effects[2], thereby significantly hampering their usage in the clinic[3]. Hence, there is an unmet need to identify novel chemical compounds with less adverse effects to combat this devastating disease.

Apoptosis is a type of cell death that is characterized by the preservation of plasma membrane integrity, which prevents local inflammatory reactions and tissue damage[4]. Both intrinsic and extrinsic pathways ultimately converge through the caspase cascade [5–7]. Apoptotic cell death has attracted increasing attention for its role in modulating inhibitory activities of anti-neoplastic compounds[8]. Indeed, an increasing number of reports have demonstrated apoptosis induction as the main mechanism for multiple anticancer agents [9].

Peiminine is a natural compound that is extracted from the bulbs of *Fritillaria* thunbergii (Liliaceae family) and Bolbostemma paniculatum (Maxim) Franquet (Cucurbitaceae family), and is widely used in traditional Chinese medicine for treating several diseases, including cancer[10]. It has been reported that peiminine repressed colorectal carcinoma tumor growth by inducing autophagic cell death[11].

However, the role of peiminine on apoptosis in HCC and its underlying mechanism of action remain largely unknown. Hence, the purpose of this study was elucidate the potential molecular mechanism of apoptosis induced by peiminine.

## Materials and methods

### Chemicals and reagents

Peiminine which purity is 99.8% was purchased from Pure-one Bio Technology, C_O_., LTD. z-DEVD-fmk was purchased from Selleckchem Co., Ltd (Shanghai, China).RPMI-1640 medium, 3-[4,5-Dimethylthiazol-2-yl]-2,5-diphenyltetrazolium bromide (MTT), propidium iodide (PI), 4,6-diamidino-2-phenylindile(DAPI), dimethyl sulfoxide (DMSO), and anti-Bax, Bcl-2, procaspase-3, −8, −9, caspase-3, −8, −9, PARPi (Asp214, 89 kD), PARP_1_ (Asp214, 89 kD) cleaved and β-actin primary antibodies were from Sigma-Aldrich Chemical Co., Ltd (Shanghai, China).

### Cell culture

HepG2 cells purchased from the Cell Bank of Type Culture Collection of Chinese Academy of Sciences (Shanghai, China) were grown at 37°C in a 5% CO_2_ incubator in RPMI-1640 medium supplemented with 10% (v/v) heat-inactivated fetal bovine serum (FBS), 100 μg/ml streptomycin and 100 U/ml penicillin.

### Cell viability assessment

The MTT assay was performed using 1×10^4^ HepG2 cells /well in 96-well plates. After culturing for 12h, cells were incubated with peiminine at concentrations of 2, 4, 6, 8, 10, 12, and 14 μg/ml for 24, 48, or 72 h, respectively. Then, 20 μl of MTT solution (5 mg/ml) was added for 4 h, medium was removed, and formazan crystals were dissolved in 150 μl DMSO. Next, the absorbance was read at 570 nm, and cell viability (%) was determined as follows:

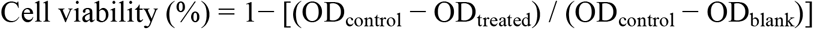

### Nuclear staining

HepG2 cells treated with peiminine at the concentration of 4 μg/ml for 24h were fixed with paraformaldehyde at 25°C for 10 min, and then incubated with 20 μl DAPI at a concentration of 10 μg/ml. Nuclear morphology was analyzed using an Olympus FV1000 fluorescence microscope.

### Detection of DNA fragmentation

A total of 3 ml cell suspension was plated into 6-well plates (1×10^6^ cells/well) and cells were treated with peiminine at concentrations of 0, 2, 4, 6 μg/ml. Cells were lysed after incubation for 24 h. Genomic DNA was extracted using phenol/chloroform/isoamyl alcohol (25: 24: 1), and RNA was removed by RNaseA. For the assay, DNA (30 μg) was separated by 1.5% agarose gel electrophoresis, visualizing DNA fragmentation patterns.

### Apoptosis assessment

After treatment of HepG2 cells with peiminine at various amounts for 24 h, apoptosis was evaluated using an Annexin V-FITC kit (BD Co., MA, Ltd, USA) according to the manufacturer’s guidelines. Analysis was performed by flow cytometry (BD Bioscience, MA, USA) using the CellQuest software (BDIS).

### Cell cycle detection

3 ml HepG2 cell suspension plated into 6-well plates (1×10^6^ cells/well) were treated with peiminine at concentrations of 0, 2, 4, 6 μg/ml and incubated in humidified incubator at 37 °C with 5% CO_2_ for 24h. Then the cells were harvested and fixed with 70% ethanol overnight, and incubated with 1 ml PI solution (20 μg/ ml in PBS with 1% Triton X-100) containing RNaseA at 37°C away from light for 30 min. Cell cycles were assessed by flow-cytometry (BD Bioscience, MA, USA) with the CellQuest software (BDIS).

### Mitochondrial membrane potential assessment

Mitochondrial membrane potential (MMP) was assessed using the Rhodamine 123 staining approach. 500 μl HepG2 cells (1×10^6^) were incubated with different amounts of peiminine for 24 h. 10 μl Rhodamine 123 dye (5 g/ ml) was added to cell suspension and cells were incubated in humidified incubator at 37 °C with 5% CO2 for 30 min. Then the cells were harvested and washed with PBS for 2 times. The fluorescence intensity and cell number were analyzed by flow-cytometry (BD Bioscience, MA, USA). The change of fluorescence intensity indicates the change of mitochondrial membrane potential.

### Determination of caspase activity

After treatment of HepG2 cells with peiminine at concentration of 4 μg/ml or a combination of peiminineat concentration of 4 μg/ml and z-DEVD-fmk at concentration of 10 μmol/L, caspase-3 activities were assessed using a specific colorimetric kit (R&D Systems, Minneapolis, MA, USA) following the manufacturer’s instructions. Absorbance was read by spectrophotometry at wavelenghth of 405 nm and caspase activity of caspase-3 was determined.

### Intracellular signaling array

Cell lysates were prepared as described above and total proteins were extracted. Intracellular signaling molecules were determined using a PathScan intracellular signaling array kit (Cell Signaling Technology, MA, USA) according to the manufacturer’s instructions.

### Western blotting

After treatment of HepG2 cells (1×10^6^) with 0, 2, 4 and 6 μg/ml of peiminine f or 24h, cell lysates were prepared and centrifuged at 12,000×g for 15 min at 4°C. N ext, the total protein content was extracted from the resulting supernatant and the prot ein concentration was quantified using the bicinchoninic acid(BCA) assay. Equal amounts (30 μg) of protein were separated by 10% SDS-polyacrylamide gel, and transferred onto PVDF membranes. After blocking with TBST containing 5% skimmed milk f or 1h, membranes were incubated overnight at 4°C with rabbit monoclonal anti-human β-actin, procaspase-3, p53, procaspase-8, procaspase-9, Bcl-2, and Bax primary antibod ies, and rabbit polyclonal anti-PARP(Asp214, 89 kD), casepase-3, casepase-8, casepase −9, and cleaved PARP(Asp214, 89 kD) primary antibodies. Then, membranes were washed PBS and incubated with horseradish peroxidase (HRP)-conjugated secondary antibodies at 25 °C for 1h. Proteins were visualized using an enhanced chemiluminescence (ECL) kit (Millipore, MA, USA).

### Statistical analysis

Data were presented as mean ± standard deviations (SD) from at least three independent experiments. Statistical analysis was performed by One-way ANOVA. *p* <0.05 was considered statistically significant.Software SPSS 17.0 was used for statistical analysis.

## Results

### Cytotoxicity of peiminine on HepG2 cells

The cytotoxicity of peiminine on HepG2 cells that were treated with peiminine at various concentrations was assessed by cellular morphology transformations and the MTT method. The results indicated that the cell population treated with peiminine was significantly reduced and exhibited dysmorphic features (Fig. 1A). In addition, peiminine treatment resulted in a time- and dose-dependent reduction of HepG2 cell viability. IC50 values of peiminine at 24-, 48-, and 72 h were 4.58 μg/ml, 4.05 μg/ml, and 3.79 μg/ml, respectively. (Fig. 1B)

**Figure 1.**
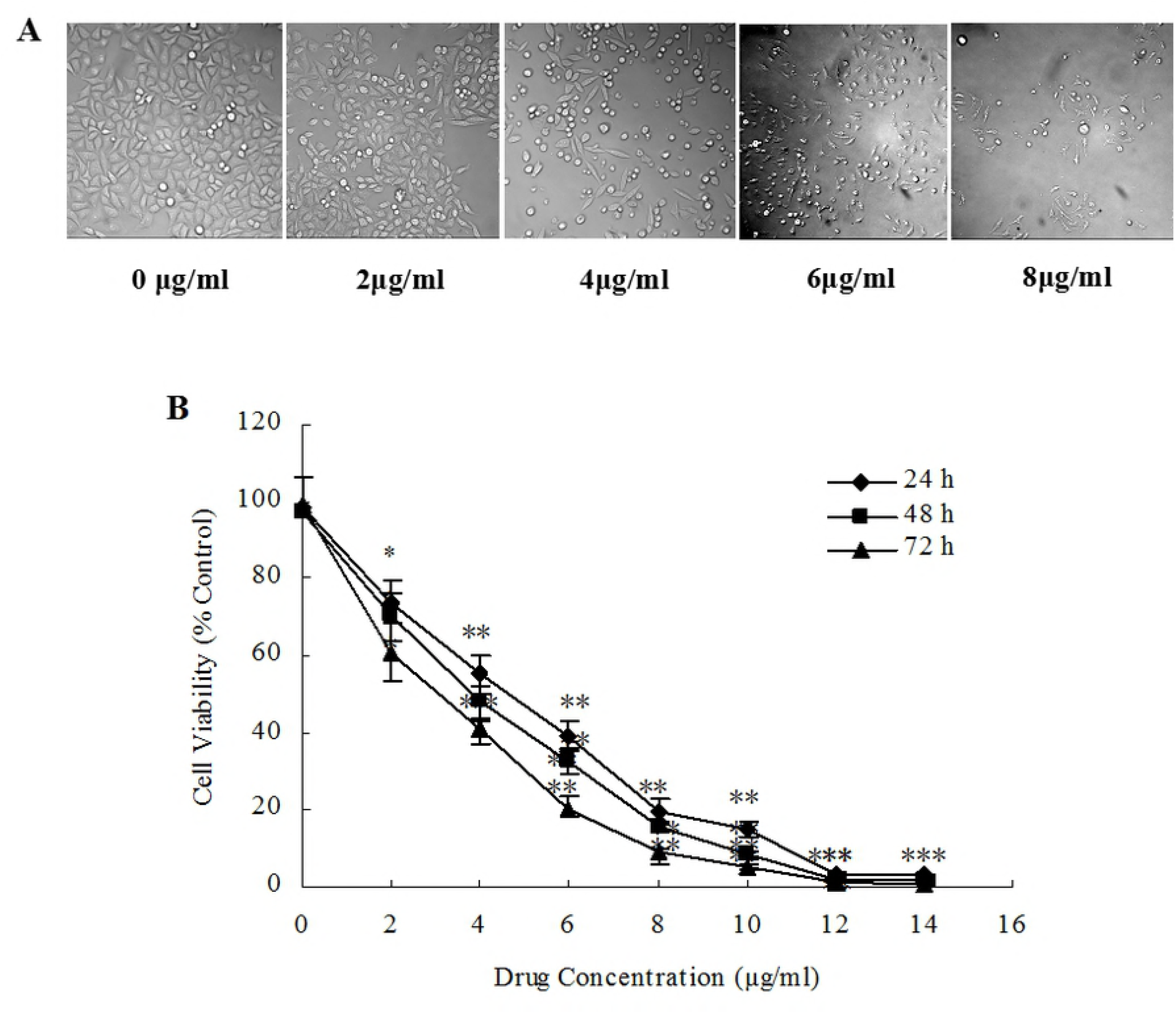
Cytotoxicity of peiminine on HepG2 cells. (A) HepG2 Cells were incubated in the presence of peiminine at indicated concentrations for 24h and cellular morphology were was using an inverted microscope. (B) HepG2 cells were incubated in the presence of peiminine at indicated concentrations for 24, 48, and 72 h and cell viability was assessed by MTT assay.

### Peiminine-induced apoptosis of HepG2 cells

After treatment of HepG2 cells with peiminine at different concentrations for 24 h, cells were stained with DAPI solution. The results showed chromatin condensation and apoptotic bodies were observed in HepG2 cells that were treated with peiminine (Fig. 2A). In addition, the result of DNA fragmentation showed peiminine can dissociated chromosome to produce DNA fragments dose-dependently(Fig. 2B). As shown in Fig. 2C and Fig. 2D, early apoptotic cells comprised 8.28, 10.32, 15.23, and 19.36 *%* and late apoptotic cells comprised 3.36, 5.08, 3.55, and 5.51% after treatment of HepG2 cells with peiminine at 0, 2, 4 and 6 μg/ml, respectively.

**Figure 2.**
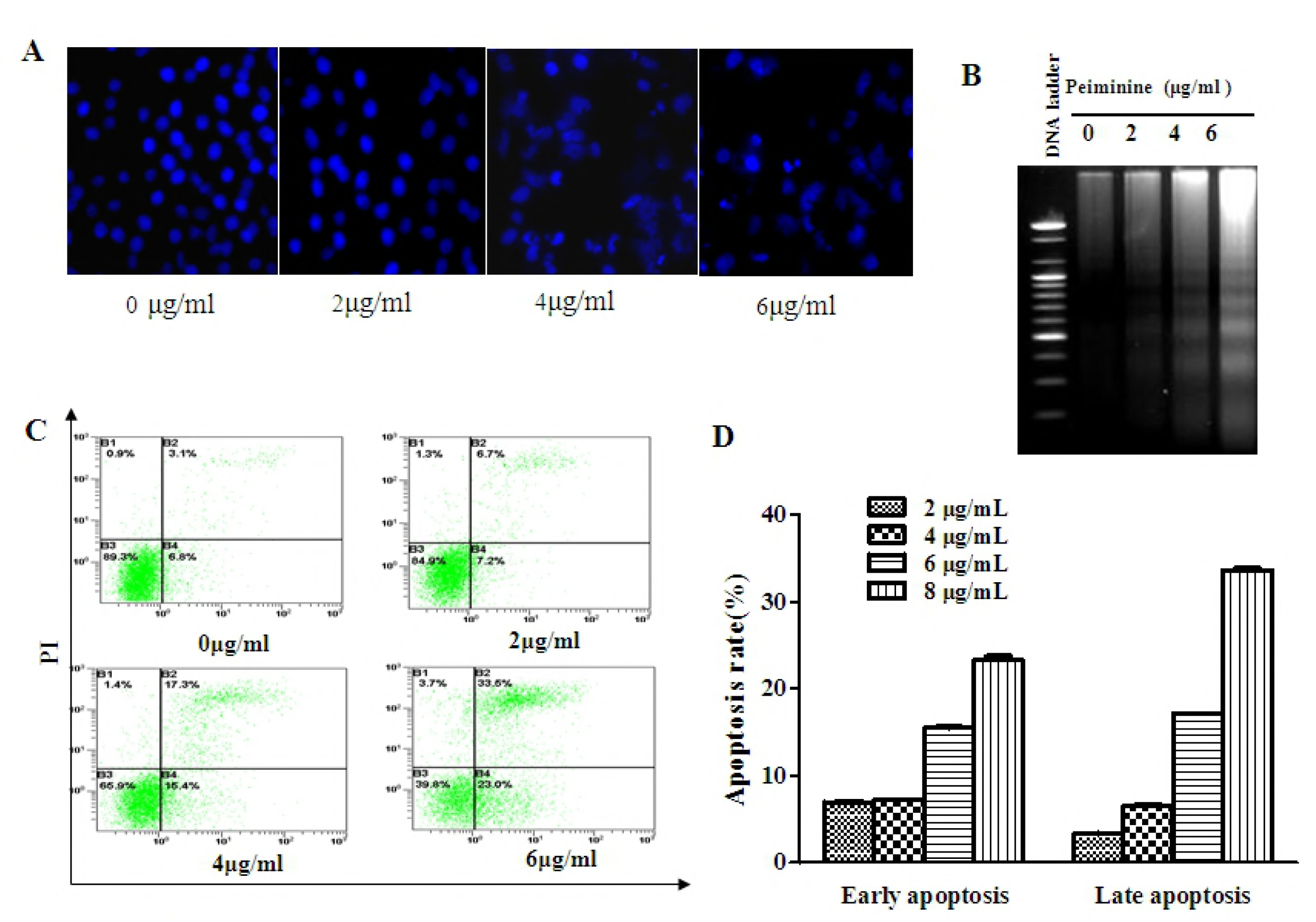
Peiminine induces apoptosis in HepG2 cells. (A) After treatment of HepG2 cells for 24 h with peiminine at indicated concentrations, cells were stained using DAPI solution. Representative images were obtained by fluorescence microscopy (×400). (B) Genomic DNA of HepG2 cells treated with peiminine for 24 h was extracted and the DNA migration profile was assessed by agarose gel electrophoresis. (C) Annexin V-FITC/propidium iodide (PI) staining of HepG2 cells after treatment with peiminine for 24 h and quantitation of apoptosis were determined by flow cytometry. (D) The apoptotic rate of HepG2 cells was analyzed by flow cytometry after peiminine treatment for 24 h.

### HepG2 cells arrest at the G2/M phase induced by peiminine

To investigate the mechanism of the growth suppression effect induced by peiminine, cell cycles distribution of HepG2 cells treated with various concentrations of peiminine were assessed by flow cytometer. The results indicated the percentage of G1 phases of HepG2 cells treated with peiminine decreased from 65.15% ±0.78 to 49.55%±0.17 with the increase of concentrations. The percentage of S phases decreased from 25.10% ±0. 52 to 11.68%±0.16 with the increase of concentrations. Meanwhile the percentage of G2/M phases increased from 17.32% ±0.20 to 39.99%±0.47 with the increase of concentrations. (Fig. 3)

**Figure 3.**
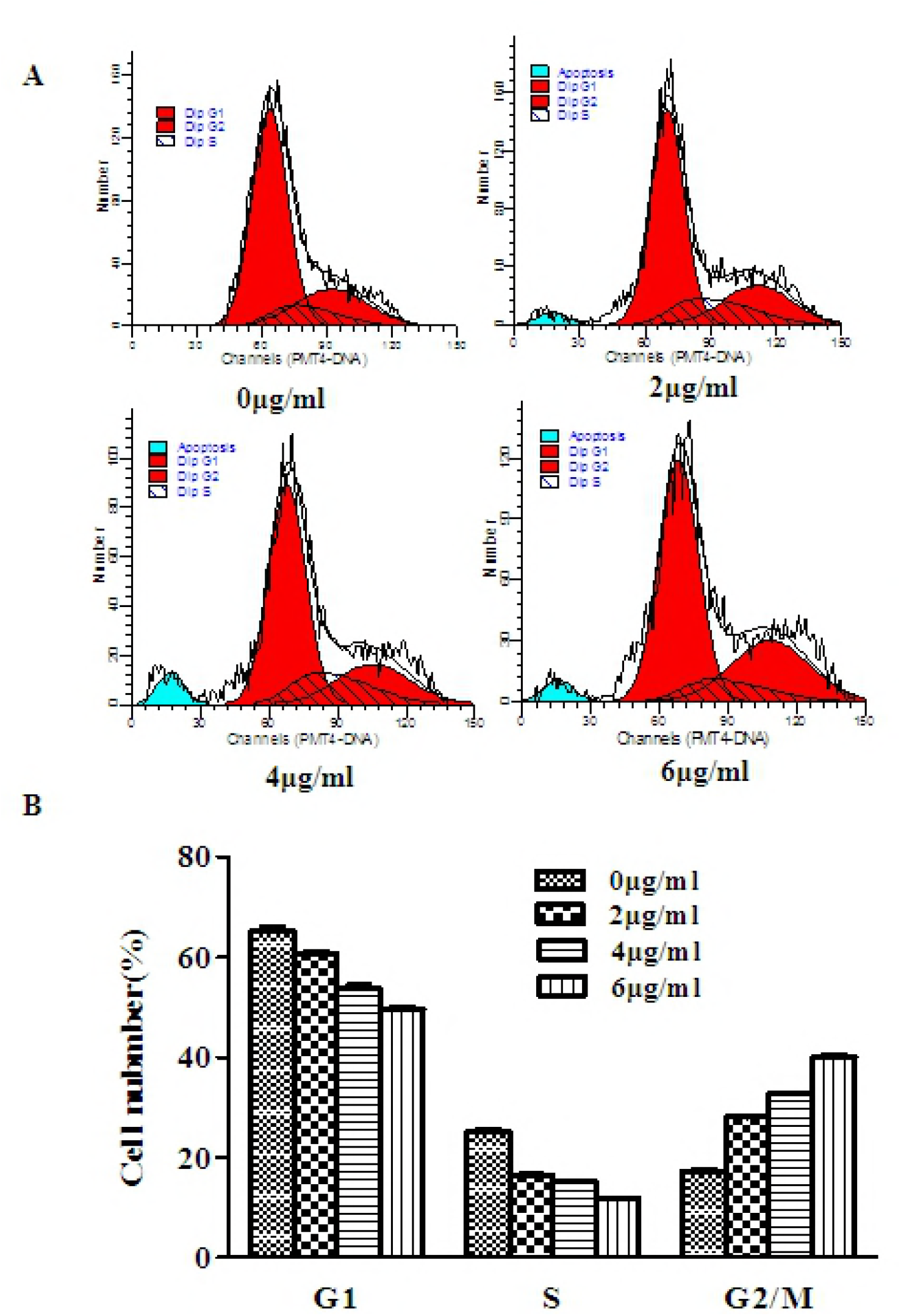
Cell cycle analysis of HepG2 cells treated with peiminine for 24 h. The values indicated the percentage of cells in the indicated phases of the cell cycle. The data shown were representative of three independent experiments.

### Mitochondrial membrane potential reduction induced by peiminine

Mitochondria activate apoptotic cell death by controlling the level of released proapoptotic proteins [12].They also have important roles in non-apoptotic cell death [13]. In this study, the results showed that the number of living cell treated with peiminine signaficantly decreased (Fig. 4A) and mitochondrial membrane potential significantly decreased in a concentration-dependent manner (Fig. 4 B).

**Figure 4.**
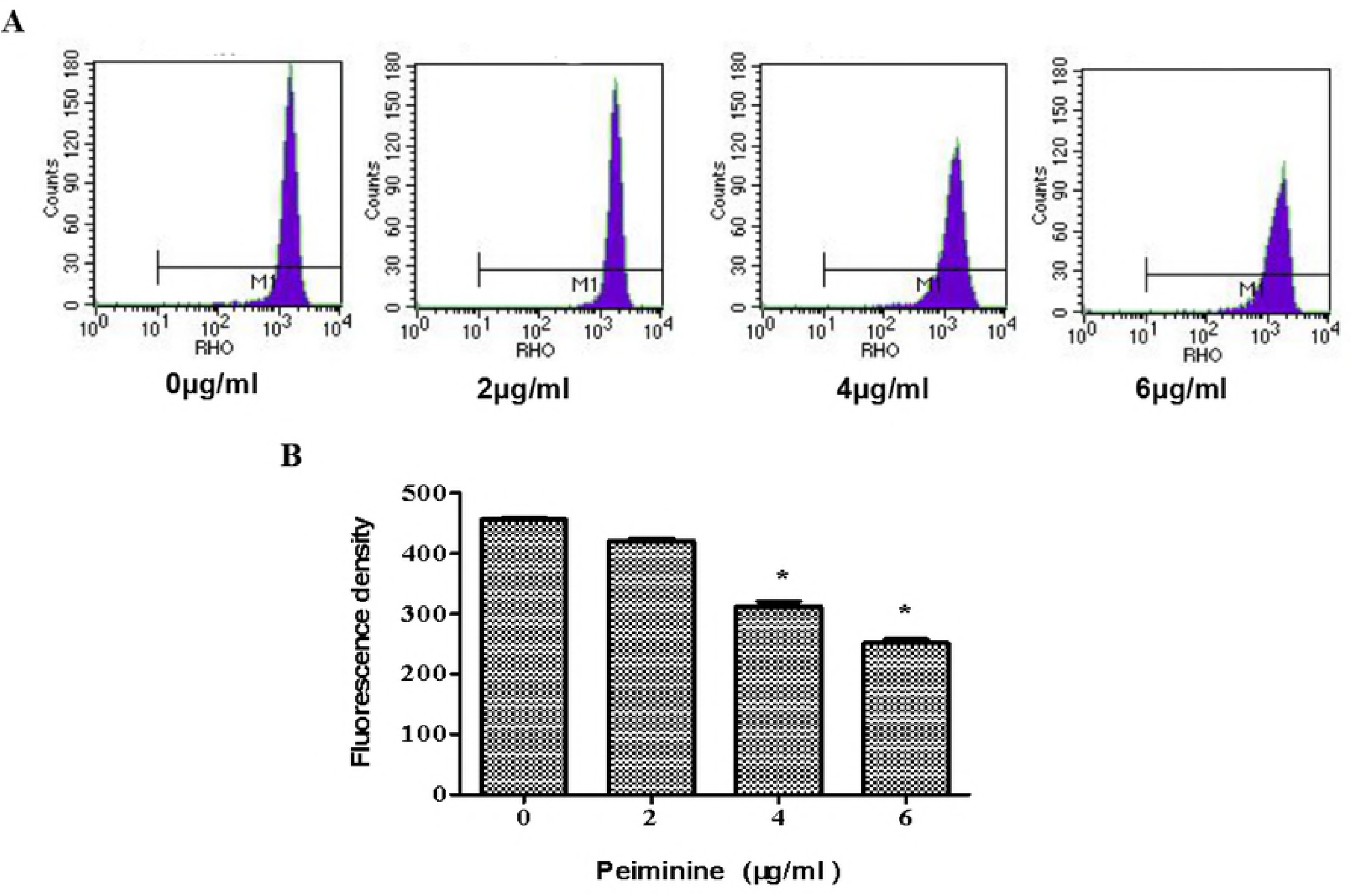
Mitochondrial membrane potential reduction induced by peiminine. After peiminine treatment at 0, 2, 4, and 6 μg/ml, respectively for 24 h, the mitochondrial transmembrane potential of HepG2 cells was determined by flow cytometry. Cells were stained using the Rhodamine 123 dye at 2.5 μg/ml for 15 min, and mean fluorescence intensity was determined by flow cytometery.

### Activity of caspase-3 promoted by peiminine

z-DEVD-fmk (Z-Asp[OMe]-Glu-[OMe]Val-Asp[OMe]-CH2F) is a cell-permeable, irreversible inhibitor of Caspase-3/CPP32[14]. In this study caspase-3 inhibitor was employed to further assess the role of caspase-3 activation in peiminine-treated cells. The results showed that z-DEVD-fmk prevented chromatin condensation of peiminine-treated cells (Fig. 5A). Moreover, DNA fragmentation also showed that z-DEVD-fmk prevented the occurrence of DNA fragmentation in peiminine-treated cells. Pretreatment of cells with z-DEVD-fmk (50 μM) markedly decreased caspase-3 induction (Fig. 5B). The activity of caspase-3 in HepG2 cells significantly increased after treatment with peiminine. However, pretreatment of cells with z-DEVD-fmk (50 μM) markedly decreased peiminine-induced caspase-3 activity (Fig. 5C).

**Figure 5.**
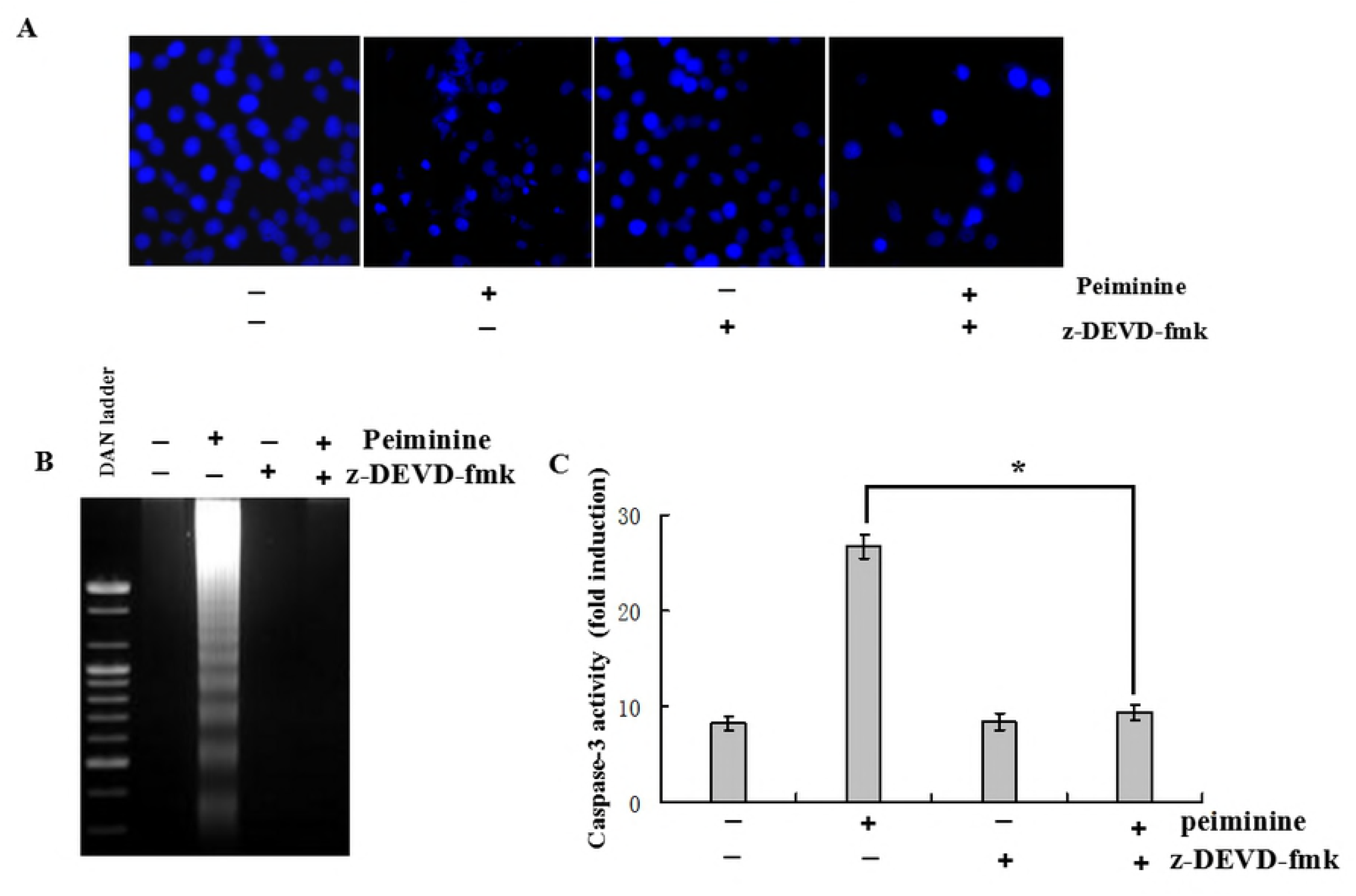
Peiminine-induced apoptosis in HepG2 cells is inhibited by a caspase-3 inhibitor. HepG2 cells were pre-incubated with z-DEVD-fmk (30 μM) for 1 h prior to treatment with peiminine (4.0 μg/ml) for 24 h. (A) DAPI staining and fluorescence microscopy analysis. (B) DNA fragmentation assessment by 0.8% agarose gel electrophoresis. (C) Evaluation of caspase-3 activity using DEVD-pNA as substrate. Data are presented as the mean±SD from three independent experiments. Student’ t-test was used for comparisons. *p < 0.05 *vs.* control (untreated cells).

### Signaling molecules of peiminine-induced apoptosis

To further elucidate the molecular mechanisms of peiminine-induced apoptosis in HepG2 cells, a PathScan intracellular signaling array kit was used to determine the changes of signaling molecules in HepG2 cells before and after peiminine treatment. The data showed that the expression of Bax, cleaved PARP, and Caspase-3,8,9 were significantly up-regulated in peiminine-treated HepG2 cells. In addition, the expression of Bcl-2 and Chk2 were down-regulated in HepG2 cells after treatment with peiminine (Fig. 6).

**Figure 6.**
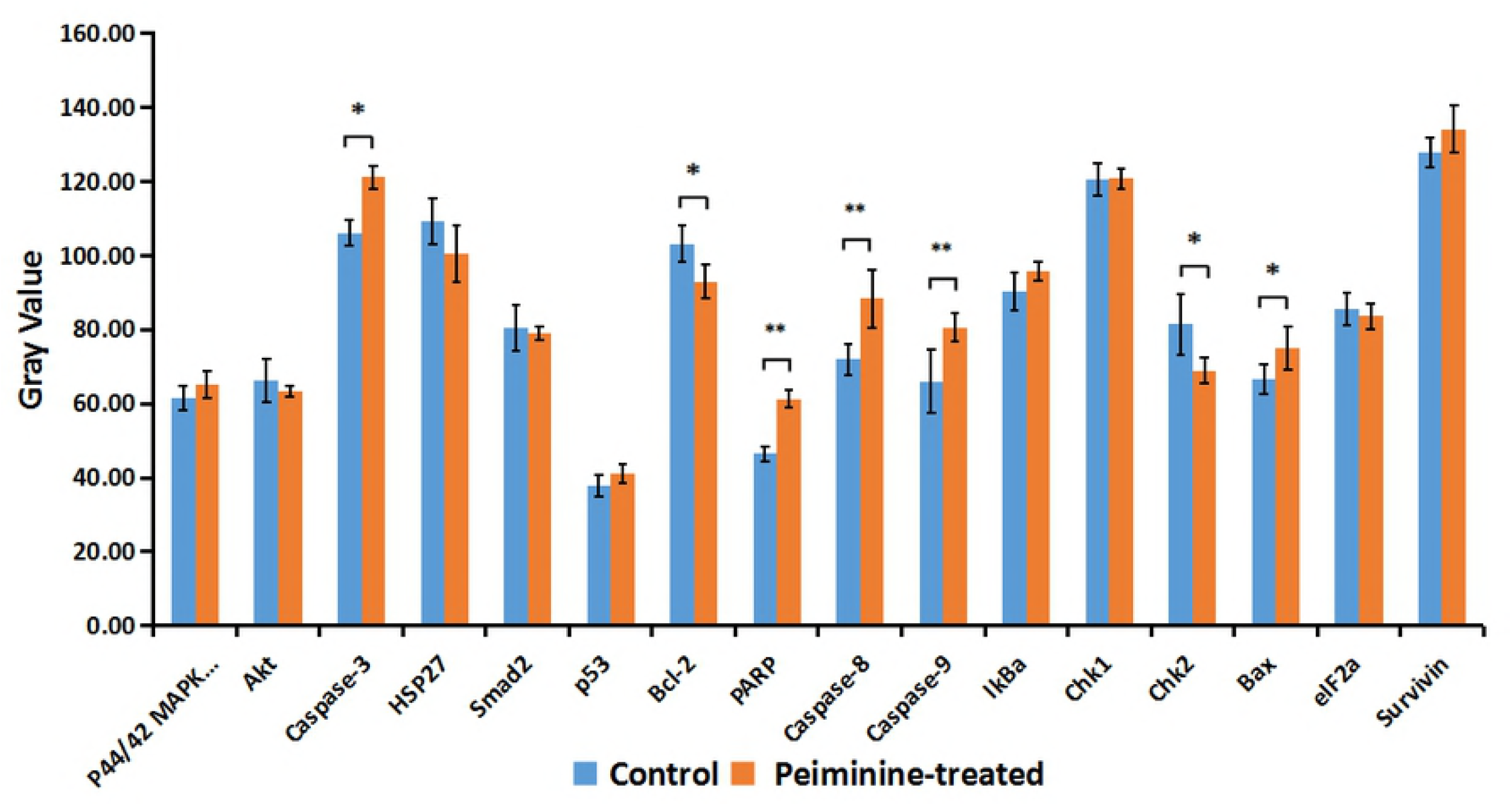
Signaling molecules of peiminine-induced apoptosis. After peiminine treatment, total proteins of HepG2 cells were extracted and intracellular signaling molecules were determined using a PathScan intracellular signaling array kit.

**Figure 7.**
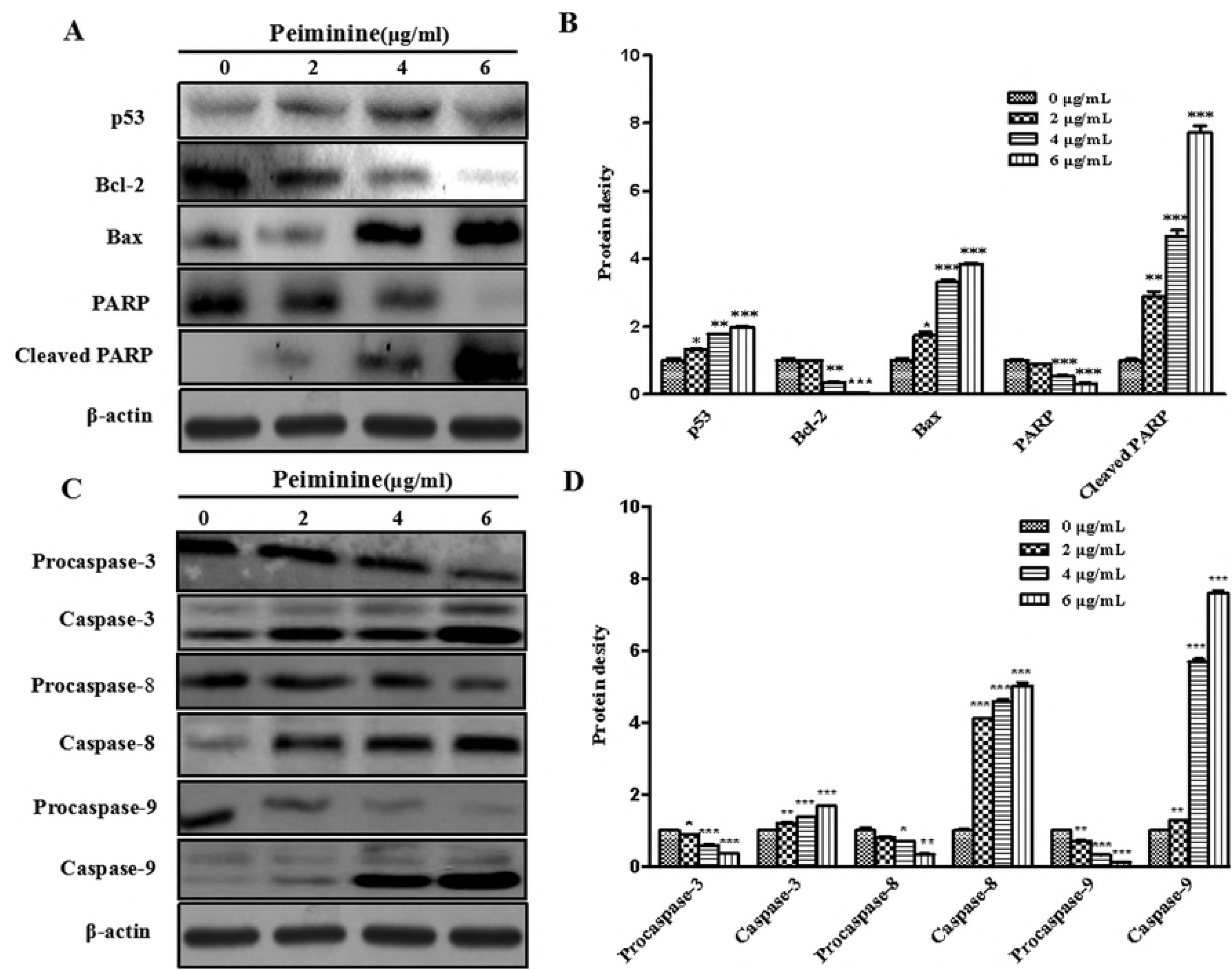
Expression of peiminine-induced proteins related to proliferation in HCC cells. (A) Expression of p53, Bcl-2, Bax, PARP, and cleaved PARP in HepG2 cells treated with peiminine were determined by Western blot analysis. (B) Expression of procaspase-3, −8, and −9, caspase-3, −8, and −9 in HepG2 cells treated with peiminine were determined by Western blot analysis. Total proteins were extracted upon incubation in presence of 0, 2, 4, and 6 μg/ml peiminine for 24 h and protein expression was assessed by Western blot analysis. β-actin served as an internal control, and experiments were performed in triplicate. *p < 0.05 vs. control (untreated cells), **p < 0.05 vs. control (untreated cells),***p < 0.05 vs. control (untreated cells)

### Effects of peiminine on the expression of apoptotic-related proteins

To further unveil the mechanism of peiminine-induced apoptosis, various key effectors of programmed cell death were quantitated at the protein level. Caspase-3 is essential for changes observed during apoptosis [15].Our results showed that the expression of procaspase-3, PARP_1_(Asp214, 89 kD), procaspases-8 and −9, and Bcl-2 were significantly reduced by 2-6 μg/mL of peiminine treatment. Conversely, caspase-3, 8, 9, PARP_1_ (Asp214, 89 kD) cleaved and Bax protein levels were significantly increased upon peiminine treatment.

## Discussion

Peiminine is a compound that is isolated from *Bolbostemma paniculatum* (Maxim) Franquet, and has demonstrated antitumor activities. In the present study, we demonstrated that peiminine markedly inhibited HepG2 cell viability in a time- and dose-dependent manner. For HepG2 cells, IC_50_ values of 4.58 μg/ml, 4.05 μg/ml, and 3.79 μg/ml were obtained at 24, 48, and 72 h, respectively. These results suggested that peiminine has potent cytotoxicity towards HepG2 cells.

Apoptosis contributed to cell homeostasis and several other physiological events [12, 13]. Two major pathways can activate caspases and apoptosis in mammalians, including the Fas /tumor necrosis factor (TNF) death receptor (extrinsic) and mitochondrial (intrinsic) pathways[15, 16] (Jin et al., 2005; Zimmermann et al., 2001). The extrinsic pathway involves caspase-8 activation, whereas the intrinsic pathway involves Bax, caspase-9, and caspase-3 [17].

Anticancer drugs often inhibit tumor cells by inducing apoptosis[18]. As shown above, morphological characteristics of apoptosis were induced by peiminine (Fig. 2). Quantitation indicated that peiminine treatment resulted in a significant increase in the number of cells in the early apoptosis phase in HepG2 cells. We examined the mitochondrial membrane potential to assess whether peiminine-induced apoptosis involved the intrinsic pathway. Indeed, peiminine treatment resulted in significantly decreased mitochondrial membrane potential.

Caspase activation results in apoptotic cell death due to death signals from cell-surface receptors, mitochondria, or the endoplasmatic reticulum (ER)[19], with caspase-3 mainly initiating apoptosis[20]. Caspase-3 activation is induced by initiator caspases, e.g. caspase-8 or −9[21].In this study, the potent caspase-3 inhibitor z-DEVD-fmk inhibited chromatin condensation in peiminine-treated cells, and attenuated caspase-3 activation.

The Bcl-2 family of proteins mainly controls the intrinsic apoptotic pathway[22], which includes pro- and anti-apoptotic effectors that are induced or inhibited according to the heterodimerization pattern[23]. In this study, we showed that peiminine treatment resulted in increased Bax levels and decreased Bcl-2 levels, thereby shifting the balance towards enhanced apoptosis. Additionally, the the expression of procaspase-3, PARP, procaspases-8 and −9, and Bcl-2 were significantly reduced by 2-6 μg/mL of peiminine treatment. Conversely, caspase-3, 8, 9, and Bax protein levels were significantly increased upon peiminine treatment.

In conclusion, peiminine displayed significant cytotoxicity in HepG2 cells and markedly increased the number of cells in the early apoptosis phase. Both extrinsic and intrinsic apoptotic pathways were triggered by peiminine, suggesting that peiminine may be considered as a potential drug candidate for liver cancer treatment.

## Acknowledgements

This study was funded by Shaanxi Provincial Health and Family Planning Commission (No.2016D027) and the National Natural Science Foundation of China (No.81774132).

## Author Contributions

**Conceptualization:** Xu Chao

**Data curation:** Guoquan Wang, Yuping Tang.

**Methodology:** Changhu Dong, Hong Li, Bin Wang, Jieqiong Wu.

**Writing-original draft:** Jiarong Zhao.

**Writing - review & editing:** Jiarong Zhao, Xu Chao

